# Accurate prediction of protein assembly structure by combining AlphaFold and symmetrical docking

**DOI:** 10.1101/2023.06.22.546069

**Authors:** Mads Jeppesen, Ingemar André

## Abstract

AlphaFold can predict the structures of monomeric and multimeric proteins with high accuracy but has a limit on the number of chains and residues it can fold. Here we show that a combination of AlphaFold and all-atom symmetric docking simulations enables highly accurate prediction of the structure of complex symmetrical assemblies. We present a method to predict the structure of complexes with cubic – tetrahedral, octahedral and icosahedral – symmetry from sequence. Focusing on proteins where AlphaFold can make confident predictions on the subunit structure, 21 cubic systems were assembled with a median TM-score of 0.99 and a DockQ score of 0.71. 15 had TM-scores of above 0.8 and were categorized as high-quality according to DockQ. The resulting models are energetically optimized and can be used for detailed studies of intermolecular interactions in higher-order symmetrical assemblies. The results demonstrate how explicit treatment of structural symmetry can significantly expand the size and complexity of AlphaFold predictions.

## Introduction

The functional complexity of cellular processes often requires the association and cooperation of multiple protein subunits in protein complexes. Protein assemblies carry out many of the cell’s most fundamental and sophisticated functions, from DNA replication to energy synthesis and molecular motion. As the molecular structure is key to understanding the function of multimeric complexes there is currently significant interest in experimental determination, driven by improvements in cryo-electron microscopy^1^, but also in the development of computational methods to predict structure from sequence^2^.

Over the last couple of years, we have seen a revolution in our abilities to predict protein structures from deep learning methods. AlphaFold (AF)^3^ and AlphaFold-Multimer (AFM)^4^ have shown unprecedented levels of accuracy in predicting structures of monomers and protein complexes and currently serve as the basis for all of the top-performing models in the latest round of the structure prediction contest CASP^5^. However, AFM is currently limited to smaller complexes as prediction accuracy decreases and memory consumption increases when more chains are modeled^6^. Fundamentally, the prediction of large multimeric assemblies presents significant additional challenges compared to that of monomeric proteins, as it requires the simultaneous prediction of the organization and interactions of multiple chains in space. The underlying training data to AF/AFM provide rich sources of information regarding the internal structure and interfaces between subunits, but less information regarding the overall organization of chains.

An attractive strategy for building larger complexes is to assemble them from predicted AF/AFM subcomponents. Recently, it was shown that large complexes with 10 to 30 chains could sometimes be predicted by sequentially assembling AFM-predicted dimeric and trimeric subcomponents by superposition^6^. However, this approach has two major challenges. First, small errors in the prediction of individual interfaces will propagate through the complexes and can result in severe clashes between subunits in the full assembly model. Second, for complexes with more than one unique interface a sequential assembly approach relies on the accurate prediction of multiple interfaces to the same subunit, which can be challenging to extract from AFM.

Moving forward it would be advantageous to develop a method that can iteratively refine all interfaces of the assembly to minimize error propagation and search for interactions not predicted by AFM using molecular docking. This has been demonstrated for heterodimeric complexes using two AF-predicted monomers and rigid-body docking methods^7, 8^. However, for large assemblies, this approach has not been explored. This is likely due to the very large number of degrees of freedom involved in the search for optimal subunit placements, leading to a computationally intractable optimization problem. Nonetheless, for multimeric assemblies displaying structural symmetry the degrees of freedom can be substantially reduced^9, 10^, making the combined AF/AFM docking approach tractable.

Many large protein complexes are either fully symmetrical, display local symmetry, or are quasi/pseudo-symmetrical (Fig. S1). Symmetry also becomes more prevalent as the protein complexes grow larger (Fig. S2). The evolution of symmetry in large protein complexes has facilitated the emergence of many shapes such as rulers, rings, and containers that are uniquely important for many functions. It is also critical for allostery, cooperativity, and multivalent binding^11–14^.

Here we present a strategy to predict the structure of large symmetrical complexes from AF or AFM subcomponents with high accuracy by combining it with an all-atom symmetrical docking method (Fig. 1A-C). We recently presented an efficient atomistic docking algorithm for heterodimeric docking called EvoDOCK^15^. In this study, we extend it to symmetry and use it to build and refine complex homomeric symmetrical assemblies built from subcomponents predicted by AFM. We demonstrate our method on a benchmark containing large protein assemblies from the most complex symmetrical systems in nature, the cubic symmetry group. The cubic symmetry group consists of the tetrahedra (T), octahedra (O), and icosahedra (I) with 12, 24, and 60 chains respectively in the simplest homomeric case (Fig. 1D-F). These complexes form spherical structures and are often involved in a variety of biological functions including the storage of genomes for many viruses. We show that our method can predict the structure of cubic symmetries from sequence at an atomic level, providing energetically optimized models of very large assemblies.

**Fig. 1:**
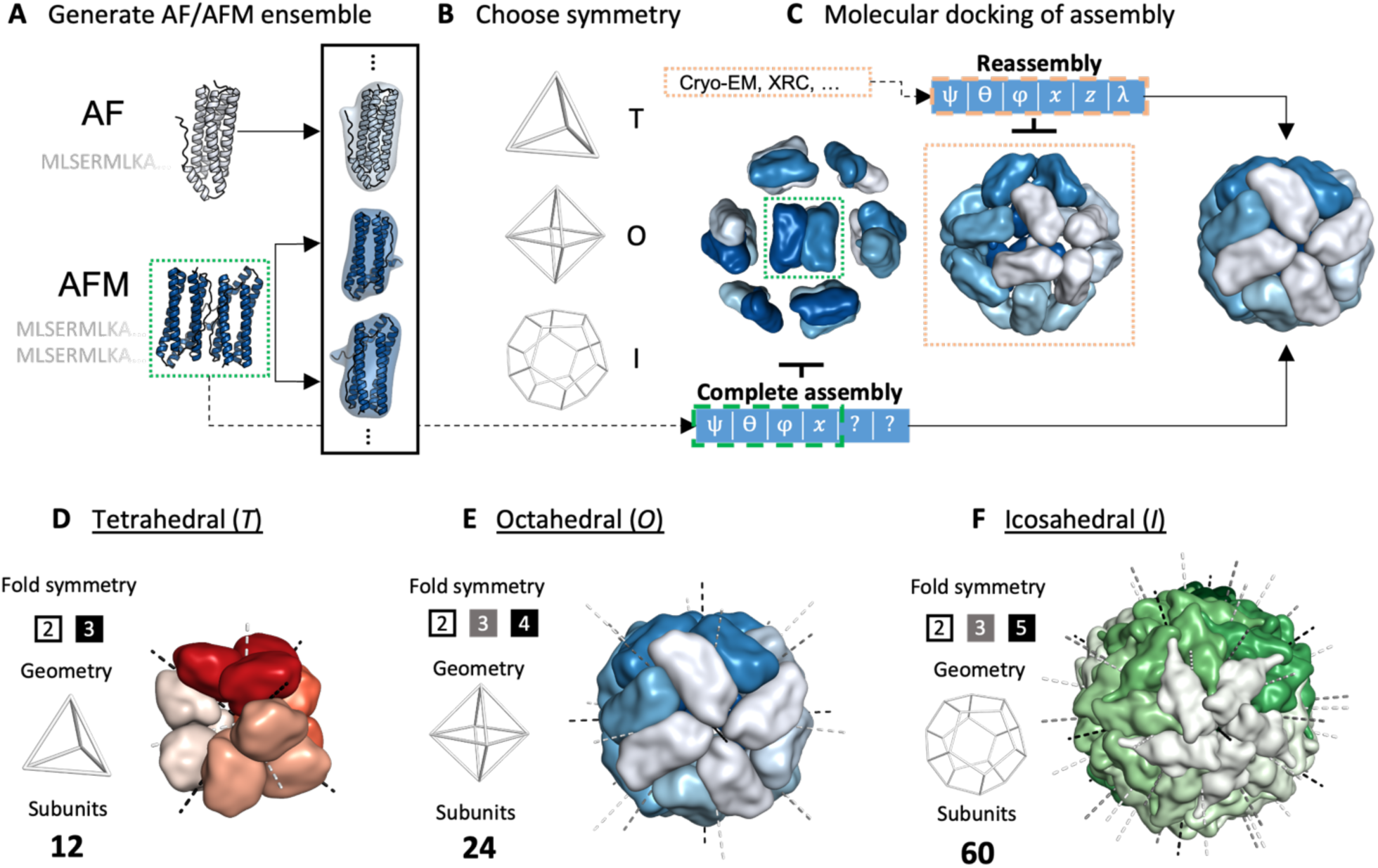
Schematic representation of the assembly prediction method and the cubic symmetry group. **A-C:** Overview of the assembly prediction method. **A:** AF (top) and/or AFM (bottom) are run on sequences of the target protein complex to produce an ensemble of input subunits. **B:** A cubic symmetric model of either a tetrahedral (T), octahedral (O), or icosahedral (I) type is chosen to model the target protein complex. **C:** The subunits are placed into the chosen symmetry and molecular docking is used to search for native rigid body parameters describing the cubic protein complex ([ψ, θ, φ, z, x, λ], see also Fig. 2B). Two approaches can be used to model the structure. With access to some structural knowledge to set the parameters, which can be derived from a variety of sources such as cryo-EM or X-ray crystallography, the structure can be reassembled (Reassembly) by searching for local optimal values of the parameters. In the absence of any structural information, the structure can be completely assembled (Complete assembly) from sequence using a combination of local docking on parameters derived from AFM and global search on the rest. **D-F:** The three cubic symmetries. **D:** Tetrahedral (T) structure having 2- and 3-fold symmetry and containing at least 12 subunits. **E:** Octahedral (O) structure having 2-, 3- and 4-fold symmetry containing at least 24 subunits. **F:** Icosahedral (I) structure having 2-, 3- and 5-fold symmetry containing at least 60 subunits.

## Results

The general methodology for predicting large protein complexes from AF/AFM subcomponents is presented schematically in Fig. 1A-C. Sequences of the target protein complex are used as inputs to AF/AFM which in turn are used to generate an ensemble of different candidate subunits for the target complex (Fig 1A). One of three cubic symmetries is chosen to model the target structure (Fig. 1B) and a symmetric version of EvoDOCK is then used to dock the assembly to produce a final energetically optimized model by optimizing the rigid body parameters describing a cubic system (Fig 1C).

Here we distinguish between two approaches to model the target structure which we call *Reassembly* and *Complete assembly* which are used in the presence and absence of initial structural information respectively. In the presence of structural information, it is often desirable to refine the corresponding model to achieve a more near-native solution. In the Reassembly approach, a local docking optimization is carried out around all rigid body parameters derived from this model (Fig. 1C, orange box). In the absence of structural information, a larger parametric space must be sampled as no templates are available. In the Complete assembly approach, the full complex is predicted directly from sequence. Through AFM, some of the rigid body parameters can be estimated and locally optimized while others must be globally optimized (Fig. 1C, green box).

A benchmark of protein structures with cubic symmetry was constructed to evaluate the method, consisting of assemblies with tetrahedral, octahedral, and icosahedral symmetries. Requirements on experimental resolution, structural diversity, and monomer size were used to select the final set. In addition, we limit ourselves to systems where AF/AFM can accurately predict monomer structures and where AFM produces some oligomers that are highly symmetrical. The size of the benchmark set was also limited to reduce computational time, since atomistic docking simulations are expensive compared to AF/AFM predictions. The final benchmark contains 21 cubic systems, 7 from each of the three symmetry groups (see Methods for the full selection procedure).

Accurate prediction of assembly structure required an extension of the EvoDOCK approach to cubic symmetry and optimization of conformational sampling for complex and large symmetrical systems, described in the next section. This is followed by a section on how to utilize AF and AFM to generate structural ensembles required for the assembly simulation. We test our methodology on increasingly more difficult scenarios. In the first scenario, we show that the native assembly structure can be recovered in Reassembly experiments using the backbone of the native monomer. In the second scenario, we show that the assembly structure can be recovered in Reassembly experiments using subcomponent structures predicted by AF/AFM. In the third scenario, we demonstrate that the structure of cubic symmetrical systems can be predicted directly from sequence without prior information on the rigid body orientation in Complete assembly experiments. Finally, as the benchmark is limited to systems that produce accurate and symmetrical AF/AFM predictions, a broader spectrum of cubic subcomponents with AF/AFM is predicted and we analyze what fraction we can expect to be successfully predicted by the method.

### Assembly of proteins with cubic symmetry using symmetrical docking

The symmetric docking step in this study is built on an extension of the EvoDOCK method^15^ to symmetry. EvoDOCK is based on a memetic algorithm that combines a differential evolution method coupled to a Monte Carlo local docking search and is presented schematically in Fig. 2A.

**Fig. 2:**
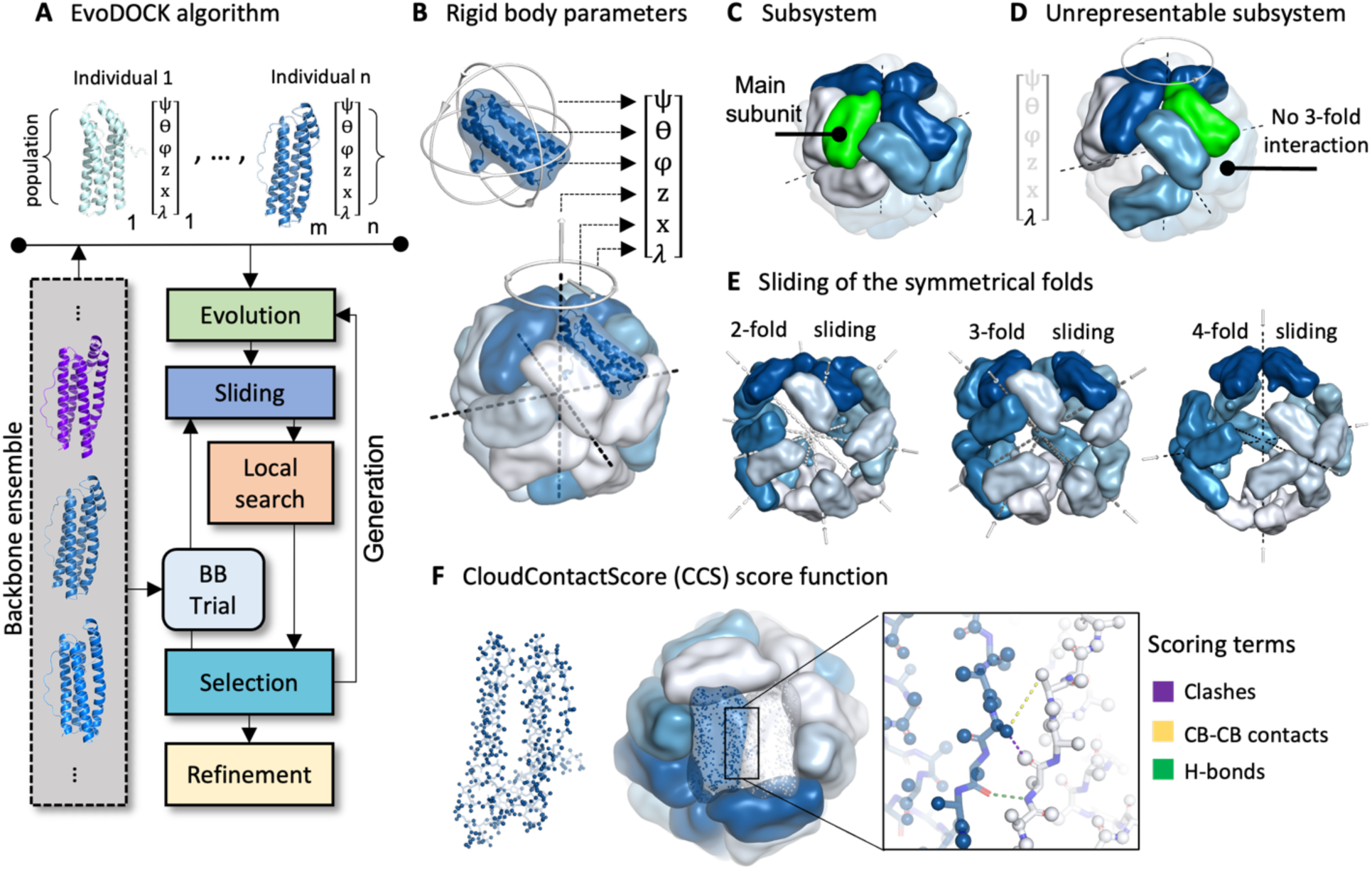
Symmetric EvoDOCK. **A**: The EvoDOCK algorithm as described in the main text. **B:** The cubic rigid body representation is composed of three parameters that control the rotation around each subunits center of mass (ψ, θ, φ), and three that control the orientation of the entire assembly (z, x, λ). **C:** The subsystem (bright surfaces) is used internally to represent the entire assembly. The subunit in green is the main subunit from which all energetic interactions are calculated. **D:** The integrity of the subsystem can be lost due to large perturbations of z, x, and λ. Here λ is perturbed beyond its bounds so that the whole structure energy will be wrong as the 3-fold interaction is not seen by the main subunit. **E:** Sliding axis of the 2-, 3- and 4-fold axis during Sliding in an octahedral case. When sliding along the 2-fold for instance, the 2-fold interface is kept fixed while the 3- and 4-fold interface contacts are improved. **F:** Left: Atoms on the surface (dark blue) of each subunit’s backbone (light blue) as represented internally in CloudContactScore (CCS). Middle/Right: Terms for the score function used in CCS to evaluate interchain interactions.

A population of individuals, each containing a randomly chosen backbone from an ensemble and six randomly chosen rigid body parameters, are initialized. The backbones and rigid body parameters of the individuals are optimized through a series of generations containing four steps: *Evolution*, *Sliding*, *Local search*, and *Selection*. During Evolution, the individuals share optimal parameters through mutation and recombination events. This drives the individuals towards more optimal solutions in the population but can introduce suboptimal energies. In the *Sliding* and *Local search* step, energies are refined by first sliding the subunits towards each other followed by an optimization of the rigid body parameters using a Monte Carlo and minimization search strategy. In the selection stage, each optimized individual is compared to its predecessor and is either continued or reverted by selecting the one with the best energy. Occasionally a new backbone is inserted into the individual from the ensemble (*BB trial*) and the *Sliding*, *Local search,* and *Selection* stage is repeated. After several generations, the whole structure is energy refined where the backbone and sidechains are simultaneously optimized and a final model is output. The optimization is guided by the all-atom energy function of Rosetta^16, 17^ and uses the symmetry machinery of Rosetta to model the structure^10^.

To adapt EvoDOCK for cubic symmetry we changed the six rigid body parameters based on a heterodimeric docking scenario to six based on cubic systems (Fig. 2B). Conformational sampling in cubic symmetry is complex due to the high packing density of subunits, the high degree of shape complementarity between chains, the presence of multiple protein-protein interfaces in the assembly and the fact that small changes in parameters can lead to drastic changes in overall assembly structure. To address these issues several improvements to the EvoDOCK methodology were made. For computational efficiency, the complete assembly structure is not modeled but rather a subsystem of chains that contains all the interactions required to calculate the energetics of the complete system (Fig 2C). To maintain the integrity of this subsystem and enable the calculation of the whole structure energy, rigid body parameters are not sampled freely but constrained within an interval (Fig 2D, Methods). This also has the benefit of making conformational sampling more efficient. During the simulation, one of the types of interfaces (2, 3, 4- or 5-fold) present in the cubic assembly may be correctly identified, while others are suboptimal. We implemented a sampling approach where the optimal interfaces can be kept, while primarily improving contacts to the other interfaces using optimized *Sliding* moves (Fig. 2E, Methods).

A fundamental challenge with sampling in cubic symmetry is to identify clash-free and energetically realistic subunit packings. This is particularly difficult using an all-atom energy representation, which produces very complex energy landscapes where tiny changes in orientation can lead to large atomic clashes. To address this problem, we designed a subroutine during the *Local search* to identify clash-free packing orientations that can be further improved in the all-atom optimization step. The subroutine does multiple rigid body perturbations guided by a score function which we call CloudContactScore (*CCS*) (Fig. 2F, Methods). Internally in *CCS*, the assembly is represented as a cloud of points consisting only of the surface atoms of the backbone of the subunits. This fast-to-calculate representation is used to quickly guide structures towards fewer clashes, more interface h-bonds, and better backbone contacts.

To test the proficiency of the methodology a Reassembly experiment was carried out for all the systems in the benchmark. The 6 parameters describing the rigid body orientations of the subunits in the assembly were randomly sampled uniformly in a broad range (Fig. S3A, Methods) centered around the values found in the native system with an average RMSD of 11Å. 100 independent EvoDOCK simulations with a population size of 100 each were started from these configurations and run for 50 iterations. In 19 out of 21 systems the lowest energy model after reassembly had an RMSD over the subsystem below 2.0 Å, while the remaining had 3.0 Å and 9.4 Å (Fig. S6). Note that the RMSD values cannot be 0 Å in all cases, because some of the experimental structures are not fully symmetrical. These results demonstrate that cubic symmetrical systems can successfully be reassembled if the native subunit structure is known. Differences compared to the native structure were primarily due to the identification of structures with alternative minima in the energy landscape, rather than inefficient rigid body sampling (Fig S4).

### Prediction of subcomponent structures from sequence using AF/AFM

Prediction of the structure of cubic systems from sequence requires prediction of subcomponent structures with AF and AFM. Predicted subcomponents are then used as inputs for the symmetric docking simulations. Using predicted subcomponents is significantly more challenging for two reasons: First, the predicted backbones will not be perfect, requiring different candidate backbones to be sampled and optimized. Second, there are parts of the structure that can only be predicted correctly in the presence of the full assembly. The N- and C-termini are often involved in the interfaces of cubic assemblies. However, AF/AFM generally cannot predict termini well and the presence of misfolded segments can prevent the formation of the correct assembly in the docking simulation. Nonetheless, a balance between removing residues with a high degree of prediction uncertainty and keeping residues that are important for interface formation must be struck.

The strategy used to produce alternative subcomponent backbones as inputs to EvoDOCK is shown schematically in Figure 3. The goal was to produce backbone ensembles with some diversity while keeping them close to the predictions by AF/AFM. We initially experimented using Rosetta^18^ to resample backbones from AF/AFM predictions as previously done in EvoDOCK^15^ and elsewhere^19, 20^ but found that backbones sampled in this manner have significantly higher RMSD to the native backbones compared to the raw output from AF/AFM. The AF/AFM predictions were therefore used directly, employing random seeds to produce different conformations. One of the strengths of AF/AFM is the ability to evaluate its prediction confidence through predicted values of the local structure quality metric LDDT^21^ (pLDDT) and the whole structure quality metric TM-score^22^ (pTM and ipTM for interfaces). These metrics were used to identify a diverse set of near-native initial backbone ensembles (Fig. 3A).

**Fig. 3:**
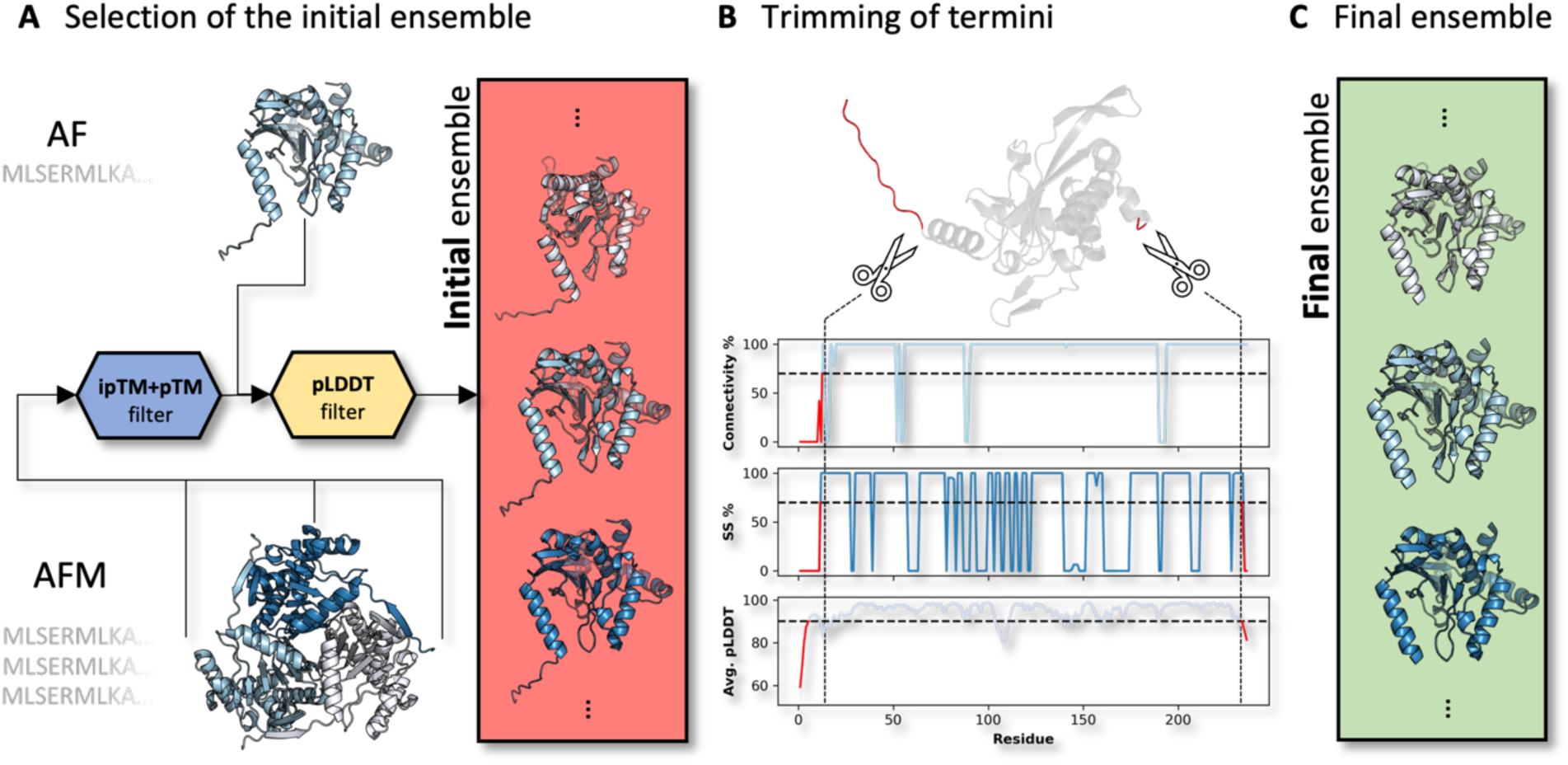
Ensemble generation strategy. **A:** AF predictions are filtered based on their pLDDT value and AFM predictions are filtered based both on their pLDDT and ipTM+pTM values to generate an initial ensemble (red box). **B:** To determine how many residues to trim at the N- and C-termini, the initial ensemble is collectively used to produce average values of three metrics per residue: connectivity (connectivity %), secondary structure propensity (SS %) and average residue pLDDT (Avg. pLDDT). Threshold values (dotted horizontal lines) for the metrics are set and terminal residues were removed from the ends until all of the metrics goes beyond their respective threshold values (indicated by the red lines). **C:** The final ensemble, which is created by removing residues as described in B, contains subunits for the target structure of equal sequence length (green box).

We utilize this initial backbone ensemble set to decide what terminal residues to remove for the docking simulations. Three metrics were used (Fig. 3B, Methods): The average residue pLDDT, the secondary structure propensity, and the residue connectivity to the rest of the structure. The pLDDT score evaluates the AF/AFM’s confidence on a per-residue basis, while the secondary structure propensity and connectivity provide measures of the expected degree of flexibility and residue-residue interaction density. Terminal residues are removed from the ends until all of the metrics goes beyond their respective threshold values (Fig. 3B, Methods). The final ensemble, which is used as inputs to the symmetric docking simulations contains subunits for the target structure of equal sequence length (Fig. 3C).

### Reassembly of proteins with cubic symmetry from predicted subunit structures

To test how well the methodology works on subcomponent structures predicted by AF/AFM we carried out the types of rigid body perturbation Reassembly experiments described previously, in which the rigid body parameters are uniformly sampled around values from a template symmetry (Fig. S3, and Methods). In our case, the parameters were calculated with the native structure as a template. However, initial rigid body parameters can also be taken from the structures of a homologous protein or estimated from a lower resolution Cryo-electron microscopy/X-ray crystallography structure in the context of model refinement.

Ensembles for subunit structures were generated as previously described using a combination of monomeric AF and AFM predictions with different oligomeric states (see Methods). The ensembles were used as inputs to 100 independent EvoDOCK simulations with a population size of 100. 100 of the best models according to the interface score (Iscore) from the EvoDOCK run were selected and refined using a symmetric energy refinement method in Rosetta^23^ (See Methods). 5 final predicted models are output by the method by clustering all 100 refined models based on the predicted rigid body parameters into 5 sets and then selecting the best model in each set according to the Iscore. The 5 models are ranked according to the Iscore and here we analyze them with respect to the best-ranked model (Best Ranked) and best model among all the clusters (Best Cluster) according to three metrics: TM-score^22^, DockQ^24^, and RMSD.

For all 5 models, we calculated the TM-score, average pairwise DockQ score across the main interfaces from all the symmetrical interfaces and the RMSD (Fig. 4A and Methods). We find median values for the Best Ranked/Cluster model as TM-score: 0.99/0.99, DockQ score: 0.76/0.82, and RMSD: 1.5/1.2 Å. We here define successful predictions as predictions having at least medium quality in their pairwise DockQ score and a TM-score of at least 0.8 and highly accurate predictions as having high quality in their pairwise DockQ score and a TM-score of at least 0.9 (Fig 4B). Under that definition, 81/86% of models are successfully predicted and 43/62% have highly accurate predictions. All energy landscapes and metric values are shown in Fig. S7 and Table S2.

**Fig. 4:**
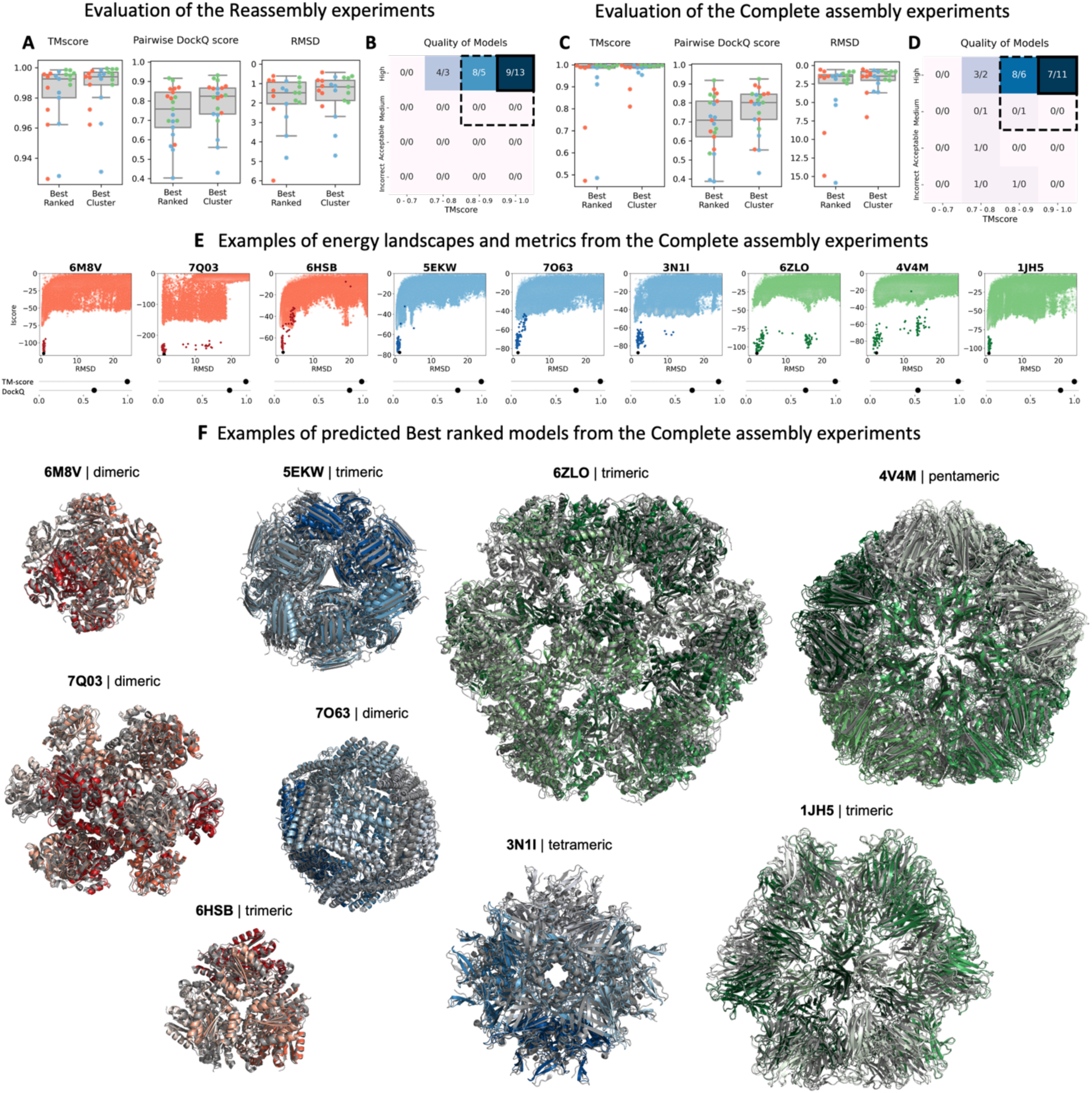
Results of the Reassembly and Complete assembly experiments. **A-B:** Results of the Reassembly experiments using subunit structures predicted by AF/AFM. **A:** TM-score, Pairwise DockQ score, and RMSD for all benchmark structures (T=red, O=blue, I=green). **B:** Classification of the results using TM-score and DockQ. The outer dashed line encapsulates successfully predicted structures and the inner highly accurate structures. **C-D:** Results of the Complete assembly experiments using single oligomeric type subunit structures predicted by AFM. **C:** TM-score, Pairwise DockQ score, and RMSD for all benchmark structures (T=red, O=blue, I=green). **D:** Classification of the results using TM-score and DockQ. The outer dashed line encapsulates successfully predicted structures and the inner highly accurate structures. **E:** Examples of energy landscapes (RMSD vs Iscore), TM-score, and pairwise DockQ score of benchmark structures of the Complete assembly experiments. **F:** Identical examples to E of the Best Ranked structures (T=red, O=blue, I=green) overlayed on their native structures in grey.

### Complete assembly prediction of the structure of cubic systems from sequence

The previous results demonstrate that with reasonable values for the rigid body parameters describing the symmetry of the cubic system, the method can successfully predict the structure of cubic complexes. In this section, we will attempt to predict the structure without any prior information or assumptions about the rigid body orientations in Complete assembly experiments. The basis of the prediction is that we can predict a single oligomeric subcomponent from AFM (dimer, trimer, tetramer, or pentamer) and use it as the starting point for the cubic assembly prediction. Ensembles of subunit structures were generated as previously described with AFM. But in addition, symmetry information was extracted from the predicted oligomer, enabling some of the 6 rigid body parameters in the docking simulation to be estimated (Fig. 2b; ψ, ϴ, φ, x, see Methods). In the EvoDOCK simulation, we sample around the individual values found in the AFM oligomer predictions. The remaining, unknown, parameters were sampled uniformly (See Fig. S3B). To get a good starting model to initiate EvoDOCK, the AFM predictions are docked along their respective symmetrical fold (Fig. 1D-F, Fig. S3B). But given that certain parameters are constrained, there are two ways to place the oligomer, related by a 180-degree flip perpendicular to the symmetry axis (Fig. S3C). In the experiments presented here, we assume knowledge of the correct orientation to spare computational resources. The end user of the method can simulate both orientations independently, or the two orientations can compete in the docking simulation. In Fig. S5 we show that EvoDOCK can learn the right orientation. In reality, it is often clear from the oligomer structure what the right orientation is. For instance, positively charged residues cluster on the inside of virus capsids to facilitate interactions with nucleic acids.

We clustered the results into 5 final models as described previously and calculated the TM-score, pairwise DockQ score, and the RMSD (Fig 4C). We find a median value for Best Ranked/Best Cluster as TM-Score: 0.99/0.99, DockQ score: 0.71/0.80, and RMSD: 1.5/1.5Å. Using the same definition for successful and highly accurate predictions as previously, it is found that 71/86% of models are successfully predicted and 33/52% have highly accurate predictions. Examples of the resulting energy landscapes from each of the cubic symmetry types for the best-ranked models are shown in Fig. 4E along with the TM-score and pairwise DockQ score (remaining systems are shown in Fig. S8 and Table S3). In Fig 4F, the predicted structures of the same examples are shown overlayed on their native structures. Taken together, the result demonstrates that accurate prediction of the structure of assemblies with cubic symmetry can be predicted with high accuracy.

### The general applicability of the method

The methodology presented so far relies on the ability of AF/AFM to produce accurate starting models for symmetric docking simulations. To get insight into the general applicability of the method, we estimated the fraction of cubic systems that can be predicted with sufficient accuracy by running AF and AFM on 111 sequences from cubic systems (see Methods). For each sequence, we attempted to predict the structure of the monomer as well as the 3 types of unique symmetric oligomers present in each cubic symmetry type (T=2/3, O=2/3/4, I=2/3/5). We define an acceptable solution as a prediction within 2 Å RMSD to the native structure. Fig. 5A shows how many structures can be predicted within this threshold for AF while Fig. 5B shows how many interfaces can be predicted for each cubic assembly by AFM.

**Fig. 5:**
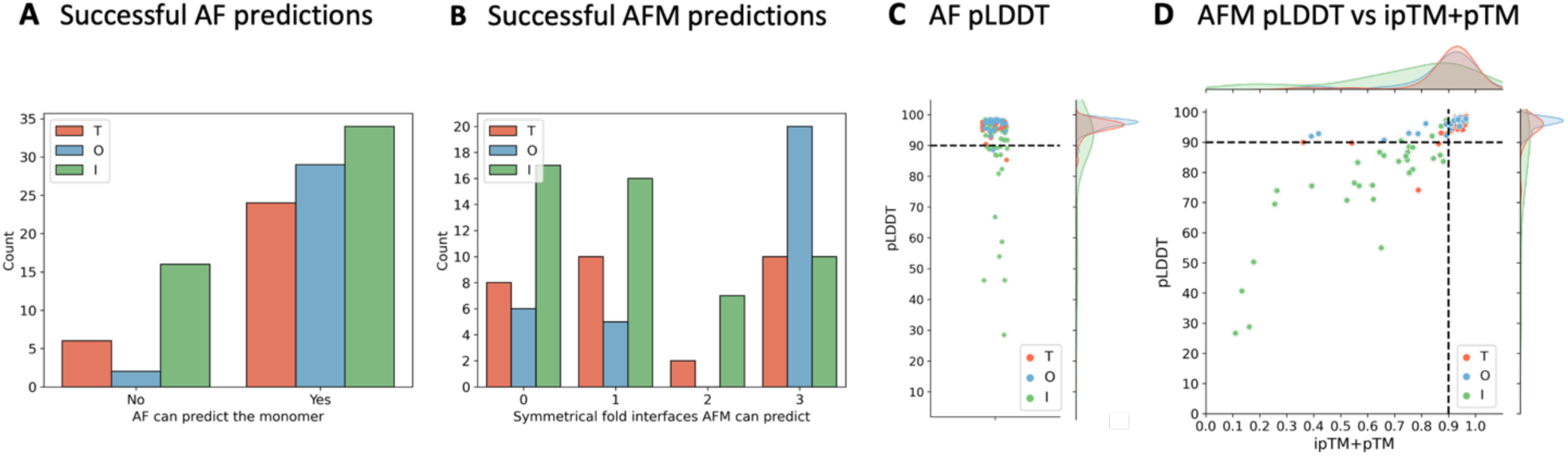
Ability of AF/AFM to predict predictions of sufficient accuracy for EvoDOCK. **A:** The number of acceptable structures AF can predict. **B:** The number of acceptable interfaces AFM can predict per assembly. **C:** Scatterplot of AF pLDDT distribution for the 111 sequences with kernel-density estimates to the right. The dashed line is drawn at pLDDT=90. **D:** Scatterplot of AFM pLDDT vs ipTM+pTM for the 111 sequences, with kernel-density estimates shown above and to the side of the figure. Dashed lines are drawn at pLDDT=90 and ipTM+pTM=0.9.

In 78% of the cases, at least one AF structure can be predicted and in 72% of the cases, at least one AFM interface can be predicted. These results demonstrate that in most cases we can expect AF and/or AFM to produce acceptable inputs for EvoDOCK. Interestingly, in 50% of the cases where at least one interface can be found, AFM cannot acceptably predict all three unique interfaces of a cubic assembly correctly. This highlights the need to use a search algorithm that can find additional interfaces outside of AFM to predict the structure of complex symmetric assemblies as we present here.

In this study, we have benchmarked only highly accurate AF/AFM predictions (>=90 pLDDT for AF and pLDDT>=90 and ipTM+pTM>=0.9 for AFM). Fig. 5C shows the distribution of pLDDT from the AF predictions. We find that 82% have this accuracy. Fig 5D shows the distribution of pLDDT and ipTM+pTM from the AFM predictions. We find that 58% of the structures have this required accuracy. For the benchmark set, we required at least one AF and one AFM prediction to be above this threshold, as well as a criterion on the symmetry of AFM-predicted oligomers (See Methods). The percentage of structures passing these filters is also 58%. It suggests that a large fraction of proteins with cubic symmetry can be predicted, even with this stringent threshold. As structure predictions evolve, this number is expected to increase. Furthermore, not limiting the AF/AFM to templates deposited before the release date of the benchmark structures as done here also suggest this number can be higher in practical cases. By combining the fraction of cubic systems with the required threshold (58%), with the acceptable solution accuracy of EvoDOCK on these systems we estimate that the current approach can successfully predict 47% or 50%, considering either the best-ranked or best cluster model respectively, of all homomeric cubic systems in a Reassembly experiment. In a Complete assembly experiment, we estimate that 41% and 50%, considering either the best-ranked or best cluster model respectively, of all homomeric cubic systems to be successfully predicted by our method.

## Discussion

Complex protein assemblies with higher-order symmetry are currently difficult to predict with deep learning methods. Symmetry puts important constraints on the structure of homomeric assemblies but is currently not directly modeled with an approach like AFM. Nonetheless, symmetry often emerges from multimer predictions of homomers so that smaller complexes can be accurately predicted. Protein complexes with cubic symmetry are far beyond what can be currently predicted with AFM due to the limit on the number of residues and chains in the current implementation^4^. The method presented here enables atomic-resolution prediction of highly complex protein assembly structures with tetrahedral, octahedral, and icosahedral symmetry by combining the capabilities of AF and AFM to model subunits and small oligomers from sequence and the capabilities of a symmetric docking protocol to model complex structural symmetry. A fundamental benefit of this approach is that it produces models that are optimized in terms of intermolecular interactions, which makes them a suitable basis for detailed analysis at the atomic level. In addition, energies provide an orthogonal quality metric that can be used to distinguish between alternative models and can be used to resolve unphysical arrangements of chains that can result from assembly approaches based on superimposition of oligomeric subsystems.

In this study, we have limited ourselves to the most complex symmetrical protein structures in nature, the cubic symmetry group. Nonetheless, the method can readily be extended for other types of symmetrical systems including those with cyclic, dihedral, and helical symmetry by optimizing against a different set of rigid body parameters, using the Rosetta symmetry machinery^9^. The approach could handle heteromeric cases as well, such as icosahedral protein capsids, by predicting heteromeric asymmetric units using AFM and using it as input for symmetric docking, although this must be tested in further benchmarking studies. We also anticipate that the same concept could be utilized to handle quasi-symmetric^11^ capsid systems with triangulation numbers higher than 1.

The method described here is limited by the accuracy of AF and AFM. We demonstrate that AF/AFM can accurately model the monomeric and oligomeric subsystems for a high fraction of cubic systems. Results in CASP15^25^ suggest that improvements in multimer predictions can be made by introducing more variation in inference by using dropouts in AFM followed by ranking by the ensembles with predicted quality metrics^26^. Such ensembles can readily be used with EvoDOCK. Our approach only requires that one of the three main interfaces in a cubic system can be predicted by AFM, and this is typically the case. This is a benefit compared to a sequential assembly approach that necessitates accurate prediction of multiple types of interfaces by AFM.

In virus capsid structures, N- and C-terminal segments are often involved in interchain interactions and may form important parts of protein-protein interfaces within the capsid. Our method does not currently model these segments. Traditional loop-modeling methods constrained by cubic symmetry could be used to complete the assembly structure. If the terminal segments reach over between different oligomeric subcomponents (domain swapping for example^27^), the current approach will fail. In that scenario, the two subcomponents would have to be modeled together in AFM. Using these more complex subsystems would require further method development.

We anticipate that the methodology presented here could be used to study cubic assemblies in several different modeling scenarios. With Reassembly experiments, the energy landscapes of native assemblies could be investigated to understand the relative importance of subunit interfaces to the overall stability of the protein. Such experiments can also be used to model the effect of mutations and to investigate assembly mechanisms of cubic assemblies. Reassembly experiments can also be used to build models of evolutionary-related assemblies, by modeling the subunit structures with AF and docking them with EvoDOCK. Another application of symmetric EvoDOCK is the refinement of structures against experimental data. EvoDOCK is implemented based on pyrosetta^28^, which can readily utilize a wide range of experimental constraints^18^, including cryo-electron densities^29^. Finally, the methodology can be used to predict structures of cubic assemblies with unknown structures. This will be particularly useful for icosahedral virus capsids. Estimates suggest that there are around 10^31^ viruses on the planet^30^, and we can hope to experimentally characterize only a fraction of this system. Nonetheless, many protein capsid proteins are substantially more complex than the homomeric systems studied here, consisting of many different types of subunits^27, 31^, having quasi/pseudo-symmetry^32^ and consisting of symmetry breaking elements^33^ and membrane anchoring. Predicting the structures of more complex biological assemblies will require more sophisticated tools than presented here but will likely require explicit treatment of symmetry and simulations of subunit assembly as we describe in this study.

## Methods

### Prediction with AlphaFold2 and AlphaFold-Multimer

For each PDB the release date in the protein data bank(PDB)^34^ was recorded. AlphaFold 2 (2.2.2) was run setting the *--max_template_date* flag to be the day before the release date of the PDB and the *--model_preset* to be either *monomer* for AF or *multimer* for AFM. AF and AFM was run as follows:

*alphafold --fasta_paths=<FASTA file path= --model_preset=<monomer/multimer> --output_dir <OUTPUT directory> ---db_preset=full_dbs --use_gpu_relax --max_template_date=<MAX template date>*

### Selection of benchmark structures

The overall selection process for the cubic structure benchmark is described in Figure 6. First a list of homomeric tetrahedral, octahedral, and icosahedral assemblies with 12, 24, and 60 chains, respectively, and with a resolution better than 4 Å was compiled from The Protein Data Bank^34^. This list was filtered based on three criteria. First, the subunit in the asymmetric unit was required to have at least one chain without chain breaks. Second, the subunit in the asymmetric unit should not include any non-canonical amino acids (except for selenomethionine, which was treated as a regular methionine). Third, the PDB file should pass through the automatic symmetry detection (see Symmetry analysis section) method to create a symmetry definition file used as a template for reassembly experiments. The remaining structures were then clustered at 90% sequence identity with CD-HIT^35^ as:

**Fig. 6:**
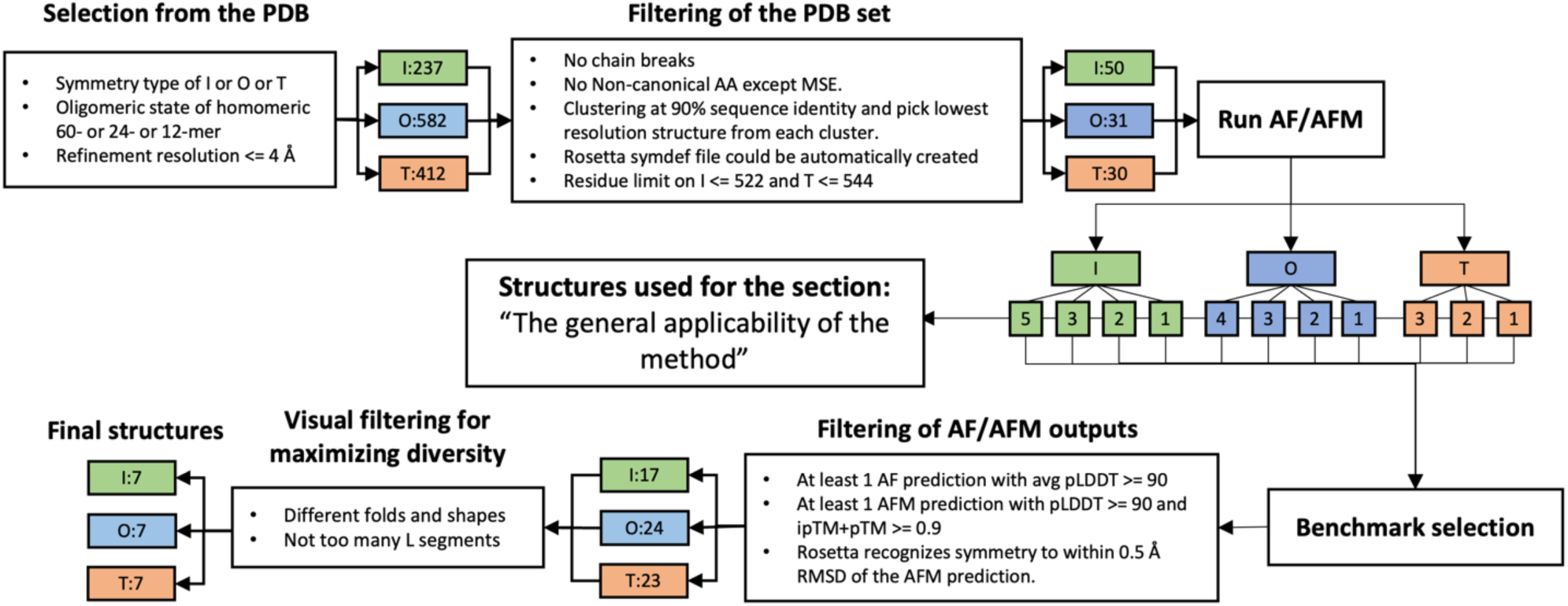
Selection of benchmark and structures for AF/AFM statistics. Information on each step is elaborated upon in the main text. After each filtering step the remaining PDBs for each cubic symmetry type (green: icosahedral (I), blue: octahedral (O), red: tetrahedral (T)) are shown. The arrows leading away from the Run AF/AFM box indicate which monomeric/oligomeric types were run for the given symmetry type.

*cd-hit -i <INPUT file> -o <OUTPUT path> -d 0 -c 0.9 -n 5 -G 1 -g 1 -b 20 -l 10 -s .0 -aL .0 -aS .0 -T 4 -M 32000*

One structure from each cluster with the best resolution was selected. The remaining structures after this filtering were then sorted based on their sequence length. To save computational time some icosahedral and tetrahedral structures were omitted if they had more than 522 or 544 residues in a single subunit respectively. This resulted in 111 proteins (I:50, O:31, T:30, Fig 6), whose sequences were predicted with AF and AFM. For AFM multiple AFM runs were launched corresponding to their symmetric fold interfaces (2-, 3- and 5-fold for I, for example). The total number of AF/AFM predictions was 414, contributing 5 models each for a total of 2070 predicted models. AF/AFM was run as described in the previous section. The generated AF/AFM predictions were used in the analysis of the fraction that pass the quality threshold for EvoDOCK assembly as described in the section: “The general applicability of the method”.

To arrive at the benchmark set, the protein systems were required to contain at least one AF prediction with an average pLDDT >= 90 and an AFM prediction (of any oligomer type) with an average pLDDT >= 90 and ipTM+pTM >= 0.9. As AFM does not enforce symmetry, only predictions where Rosetta could symmetrically reconstruct the oligomer with a CA RMSD of less than 0.5 Å to the oligomer predicted by AFM were used. Finally, manual inspection of the remaining structures was used to achieve as structurally diverse a set as possible, considering fold, shape, and loop conformations balances within each symmetry type. The final set contained 7 structures from each symmetry adding up to a total of 21 cubic structures.

### Symmetry analysis

A script was developed to automatically analyze the symmetry of native structures with cubic symmetry. The script takes the structure of the complete assembly together with the symmetry type and calculates the 6 parameters describing the degrees of freedom used in the EvoDOCK simulation. The output is a symmetry definition file used to model symmetry in the Rosetta symmetry machinery^10^. To analyze how symmetric a subcomponent within a natural cubic assembly is, we use the *make_symmdef_file.pl,* script provided by Rosetta and described in Dimaio et al.^10^

### A symmetric version of EvoDOCK

EvoDOCK for heterodimeric docking has been described previously^15^. The program was extensively modified to accommodate symmetry. This included developing 1) a new set of degrees of freedom, as presented in Figure 2B. 2) A new contact-based representation and energy function (CloudContactScore) for identifying clash-free and well-packed subunit orientations. 3) Parameter constraints to enable the use of subsystems during rigid body sampling. 4) A set of rigid body sliding moves to establish contact between subunits during the assembly process. 5) A local search strategy adapted to cubic symmetry to optimize all-atom energy in the system. These developments are described below.

### CloudContactScore (CCS) point cloud representation

To utilize the CloudContactScore score function an atomistic protein structure is turned into a cloud of points. Due to the symmetry machinery, only one subunit must be converted into this representation as symmetry expansion automatically creates all other copies in the assembly. The first step of the point cloud generation is to remove surface residue information on the surfaces beyond β-Carbon atoms, which is done in two steps. The Solvent Accessible Surface Area (SASA) is calculated across all residues and all residues with more than 20 Å SASA-value are labeled as surface residues. Then the SelectResiduesByLayer class in Rosetta, which identifies residue burial based on the number of sidechain neighbors within a cone along a vector from the α-Carbon (CA) and β-Carbon (CB), is used to determine surface residues. All identified surface residues are then changed to Alanine residues except for glycines. A final SASA calculation is carried out on this new structural representation and all atoms with 0 Å SASA are removed. Only backbone surface atoms of N, C, O, CA, and CB remains at this stage, whose coordinates are used as points in the point cloud representation.

### CloudContactScore (CCS) energy evaluation

There are four main terms in the CCS score function. First, a n_*clashes* term that penalizes according to the number of clashes. Two atoms are recorded as clashing if the Lennard Jones sphere, as defined per atom in Rosetta, overlaps by more than 20%. Nitrogen-Oxygen interactions can interact through a hydrogen bond and the clash distance is therefore reduced to 1.2 Å. β- Carbon (CB) Lennard Jones sphere-values are reduced to 1.5 Å to allow for closer interactions on the surface of the structure. To severely penalize clashes, each clash adds a large penalty to the score. Second, a backbone-backbone hydrogen bonding score that uses the *hbond_sr_bb* and *hbond_lr_bb* score terms in REF2015^17^ is used to model hydrogen bonding. They model short and large-range hydrogen bond terms, respectively. Lastly, a *n_cb_cb_interactions* term is used to pack the interaction between different chains by counting CB-CB contacts. The threshold distance for considering CB-CB interactions in the energy calculations is set to 12 Å. Each CB interaction bonus is weighted by the relative connection density of each CB to its subunit. The connection density is defined as:

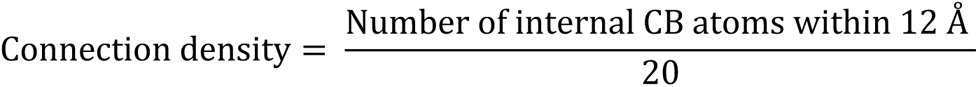

If the value of the connection density is less than zero, the CB atom is weighted 1.0. The final CCS score is given as a linear combination of the four scoring terms:

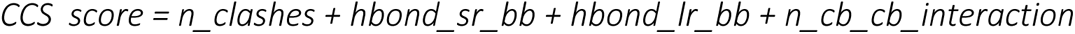

### Parametric constraints

The evasion of sampling of nonsensical rigid body orientations is achieved by bounding the rigid body parameters to ranges that maintain the integrity of the subsystem used to model the complete symmetry. For the parameters controlling the radius of the container (z) and radius of the largest n-fold symmetric system (x), they must have values above 0. A large penalty is added to the score if its parameters are sampled outside this bound using the SQUARE_WELL penalty class in Rosetta with a depth of 10^9^. High values of the λ rotation parameter can also produce nonsensical models. 4 types of symmetry input files are based on the symmetric folds of the cubic structures: 2-, 3-, 4-and 5-fold. The maximum bounds of the λ parameter are set to:

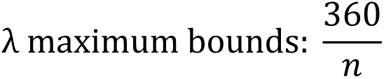

Where n is the n-fold symmetry file used (see Supplementary Table 1 for which fold symmetry was used for each structure in the benchmark). The bounds are centered around 0 and modeling icosahedral symmetry with a 2-fold symmetry input file, for instance, would yield bounds of [-90, 90] degrees. For the Reassembly experiments, half the values of the maximum bounds are used and for Complete assembly docking the full bounds are used. The other parameters have bounds of ϕ: [-40,40] degrees, ϴ: [-40,40] degrees, φ: [-40,40] degrees, and x: [-5,5] Å. These parameters are centered around the template symmetry in the Reassembly docking or the values found in the AFM predictions for Complete assembly docking. For Reassembly docking z has the bounds: [-5, 5] Å and for Complete assembly docking: [0, 1000] Å.

### Sliding moves

All subunits of the cubic system are sequentially slid along the symmetric folds from the highest to the lowest. Tetrahedral structures are slid along their 2- and 3-fold symmetry axis. Octahedral structures are slid along their 2-, 3- and 4-fold symmetry axis. Icosahedral structures are slid along their 2-, 3- and 5-fold symmetry axis. Each fold-symmetric partner is kept fixed relative to each other at each step. The sliding happens in steps of 0.3 Å and ends when clashes are detected according to the CCS *n_clashes* term or 100 sliding moves have been tried without any clashes emerging. If the structure goes out of bounds it is reverted to the starting configuration.

### Local search

The local search consists of two main components. The first part is a packing-minimization step that consists of a Rosetta-based sidechain optimization (packing) step followed by a Rosetta-based gradient minimization step that occurs if the energy was decreased by 15 units during the packing step. Both methods use the Rosetta REF2015^17^ score and a Metropolis Criterion to accept the final structure. The second part is a quick rigid body search subroutine consisting of 10 rigid body moves that use the CCS score and Metropolis Criterion to accept. Overall, the packing-minimization step occurs first, followed by the rigid body search, followed again by a final packing-minimization step.

### Ensemble generation

For the Reassembly experiments, AFM was run to produce a total of 100 single chain subunits stemming from each cubic symmetry type’s respective symmetric folds. AF was run to produce 100 chains. Thus, AFM and AF generated a total of 200 totaling subunits for each system (Fig 7A). For the Complete assembly experiments, AFM was run to produce a total of 200 chains stemming only from a single oligomeric subcomponent (Fig 7A).

**Fig. 7:**
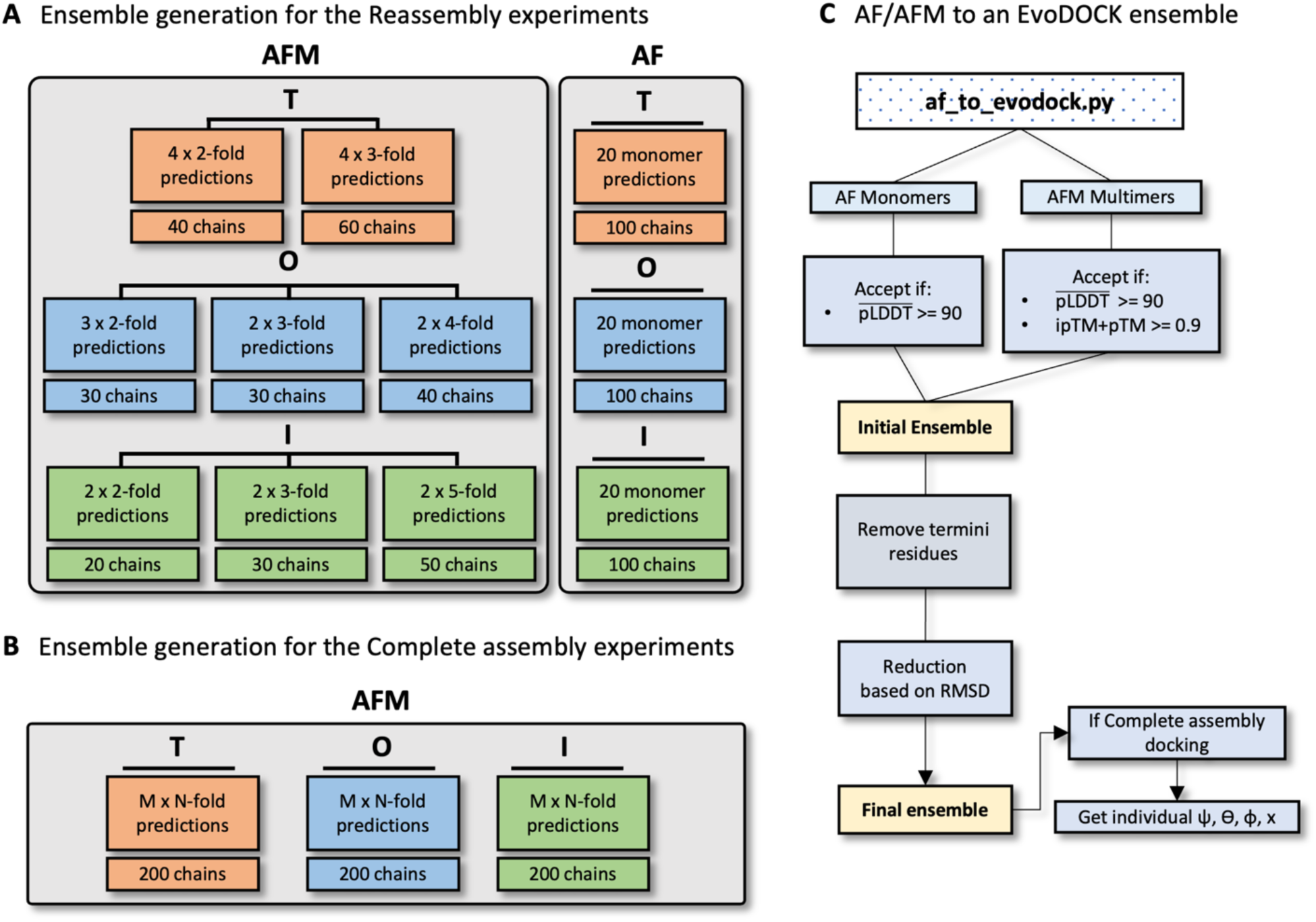
Ensemble generation methodology. **A:** Ensemble generation strategy for the Reassembly experiments. Each box for AFM (left) and AF (right) shows how many predictions for each cubic symmetry type were run and how many chains they produced in total. **B:** Ensemble generation for the Complete assembly experiments. For a given oligomeric prediction (N-fold) AFM is run M times to produce a total of 200 chains. For instance, for a 5-fold prediction M is 40 as AFM produces 5 chains for each prediction. In case of the 3-fold, more predictions were run and finally only 200 chains selected. **C:** The internals of the af_to_evodock.py script that is run to generate an input ensemble for EvoDOCK from AF/AFM predictions. Boxes are described in the main text.

AF was run with the flags as described previously, multiple times in succession, to achieve the number of target predictions. AFM was run with the flags as described previously but including: *--num_multimer_predictions_per_model* to achieve the requested number of target predictions.

The methods internal script used to generate the ensemble from AF/AFM predicted subunits (encoded in the script *af_to_evodock.py)* is shown in Figure 7C. All AF monomer predictions were first filtered based on their pLDDT (>= 90 for the benchmark structures) and all AFM multimer predictions by their pLDDT and ipTM+pTM scores (>= 90 and >= 0.9 respectively for the benchmark structures) to produce a final ensemble (as in Fig 3C). The termini were then removed as described in Fig. 3B from the remaining subunits in a process described here in further detail, using pLDDT, secondary structure propensity and residue connectivity. The average residue pLDDT is taken directly from the AF/AFM output. The secondary structure propensity is calculated with the help of DSSP^36^ while the residue connectivity was calculated with a custom function in the *af_to_evodock.py* script. For this, a contact map is created, and a residue is designated ‘disconnected’ if it is not in contact (8 Å) with another residue 10 residue neighbors downstream/upstream to the rest of the structure. These three metrics are collectively used to determine which residues to remove as described in the main text.

The ensemble is further reduced to remove structural redundancy by iteratively removing very similar structures in the ensemble. All pairwise RMSD values are calculated and members with values below 0.1 Å are kept, using the model with the best prediction metrics. However, if the final size of the ensemble is less than 50 predictions, the threshold for similarity is reduced by lowering it 0.005 Å for up to 18 steps.

Finally, a set of models for the ensemble is generated. In the Complete assembly experiments, we extract starting values for some rigid body parameters (ψ, ϴ, φ, and x) from the AFM oligomer predictions, and sample around those.

### EvoDOCK simulations

Symmetric EvoDOCK was run 100 independent times with a population size of 100 individuals. For the Reassembly experiments with a fixed backbone, 50 iterations were run while in the other simulations, 100 iterations were used. The mutation rate was set to 0.1, while a value of the recombination parameter of 0.7 was found to be optimal. For the Reassembly and Complete assembly experiments the rigid body parameters were initially uniformly sampled within their bounds as described in the parametric constraints section. For the Complete assembly experiments the initial z parameter was however determined by sliding the subunits away and then onto each other again using the CCS score function to stop the sliding when clashes were detected. The template symmetry for the Reassembly experiment is derived from the target native structure, while for the Complete assembly experiments an ideal symmetry is used. Differently from the previous scoring implementation of heterodimeric EvoDOCK is the use of the interface energy (Iscore). We noticed that interactions within a subunit could bias the selection process without improving the overall assembly energy and therefore the Iscore is used as the selection criteria.

### Symmetric energy refinement

1000 of the best structures based on the interface score (Iscore) were selected from the EvoDOCK runs and k-means clustering from the sci-kit-learn python package^37^ was used to put models into 100 clusters based on their final 6 rigid body parameters. The best models according to the Iscore values within each cluster were selected, to produce a final set of 100 models as inputs to the Rosetta FastRelax method^23^.

### Clustering of models

The 100 energy-refined structures were put into 5 clusters based on their 6 rigid body parameters using k-means clustering from the sci-kit-learn python package^37^. One model from each cluster was selected based on their Iscore resulting in 5 total models. The 5 models are the ones used to evaluate the TM-score^22^, pairwise DockQ^24^ score, and RMSD.

### TM-score

The TM-score^22, 38^ was calculated as follows:

*MMalign <INPUT file> <NATIVE file> -ter 0*

With both the input_file and native_file being the full biological assembly.

### Pairwise DockQ score

The pairwise DockQ score^24^ was calculated by summing up the DockQ score for each unique interface respective of each cubic symmetry type: 2-, 3-fold for T; 2-, 3-, 4-fold for O; 2-, 3-, 5-fold for I. For each fold, two chains that form part of the unique interface that matches best according to the RMSD to the two chains of the experimental structure were used in the DockQ score calculation. The two chains were first aligned by their residue numbers as follows:

*./DockQ/scripts/fix_numbering.pl <PREDICTED dimeric interface> <NATIVE dimeric interface>*

The output of fix_numbering.pl was then used to calculate the DockQ score as:

*python DockQ.py <ALIGNED predicted dimeric interface> <NATIVE dimeric interface>*

The pairwise DockQ score was then calculated by summing up the individual dimeric DockQ scores and normalizing them by their ΔSASA (Change in SASA when moving the chains away and back into their original position) as follows:

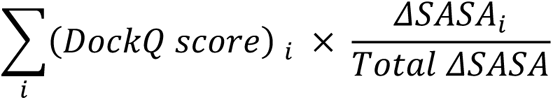

### RMSD

RMSD was calculated using the Rosetta software^18, 28^. The RMSD was calculated on the subsystem as described in the main text compared to the experimental structure. The number of different chain combinations to compare on each evaluation for the full structure is computationally intractable and the subsystem is therefore used. To make sure the full symmetrical system is captured in the RMSD calculation, each symmetrically equivalent configuration of the subsystem is used to calculate the RMSD. This is achieved by rotating around one of the symmetric n-folds n times. For instance, for an icosahedral structure, 5 configurations are tried by rotating the structure around the 5-fold separated by 72 degrees. So RMSD is calculated at 0, 72, 144, 216, and 288 degrees. The lowest value of the RMSD is selected as the report RMSD.

## Data Availability Statement

The code is available for download at https://github.com/Andre-lab/evodock, while the data and scripts for the manuscript are deposited at https://github.com/Andre-lab/af_cubic_docking.

## Author Contributions Statement

Mads Jeppesen: Conceptualization, Methodology, Software, Formal analysis, Investigation, Writing of original draft and editing.

Ingemar André: Conceptualization, Methodology, Software, Resources, Supervision, Writing of original draft and editing.

## Additional Information

The authors declare that they have no conflict of interest.

This work was supported by the European Research Council (ERC) under the European Union’s Horizon 2020 research and innovation program [771820]

