## Supplementary information for "Accurate prediction of protein assembly structure by combining AlphaFold and symmetrical docking"

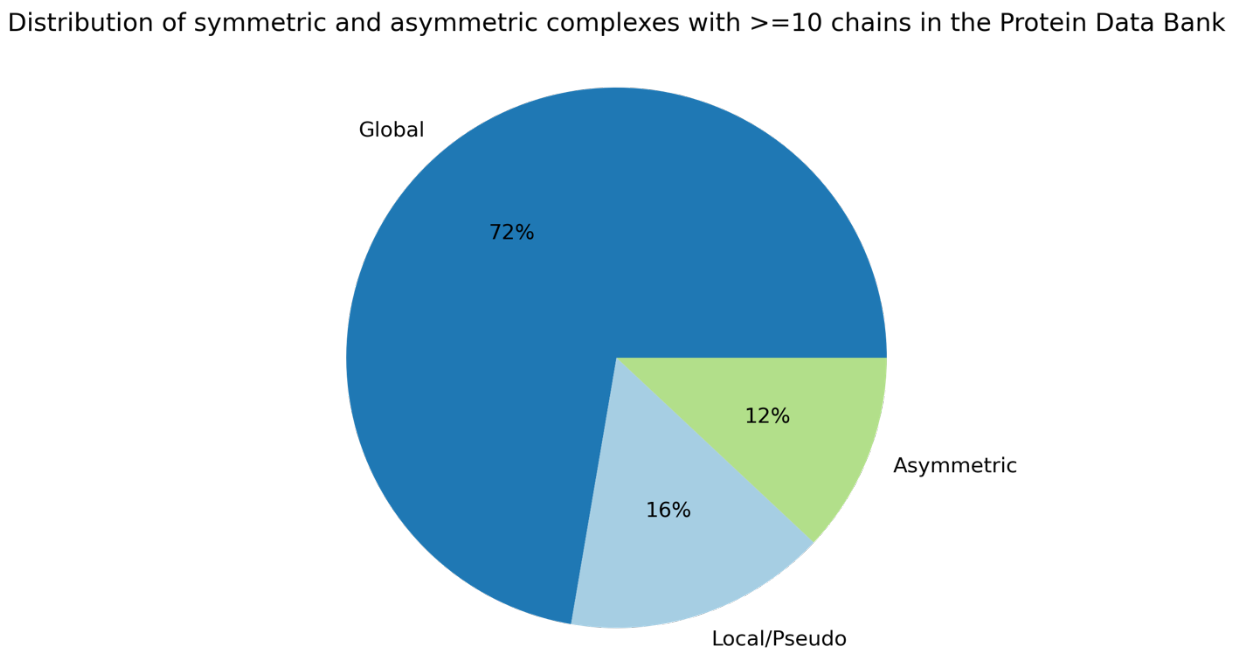


**Fig. S1: Distribution of symmetric/asymmetric complexes with more than 10 chains in the Protein Data Bank**. The Protein Data Bank[1] was culled of all PDBs containing 10 chains or more and containing no nucleic acids. These were then reduced by 30% sequence identity to a total number of 2062 individual PDBs.

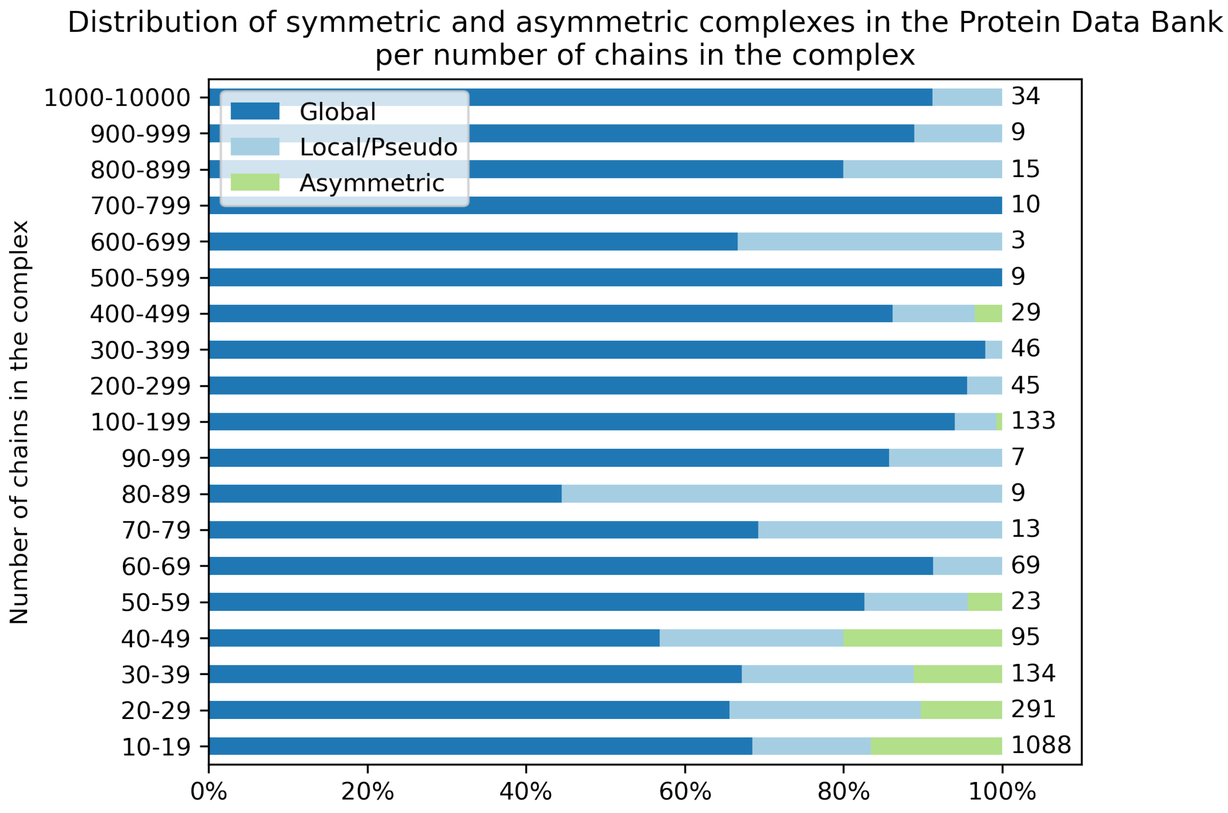


**Fig. S2: Distribution of symmetric/asymmetric complexes in the Protein Data Bank per number of chains in the complex.** PDBs were culled as in S1 and binned from the number of chains in their complex (left y-label). The total amount in each bin is shown in the right y-label.

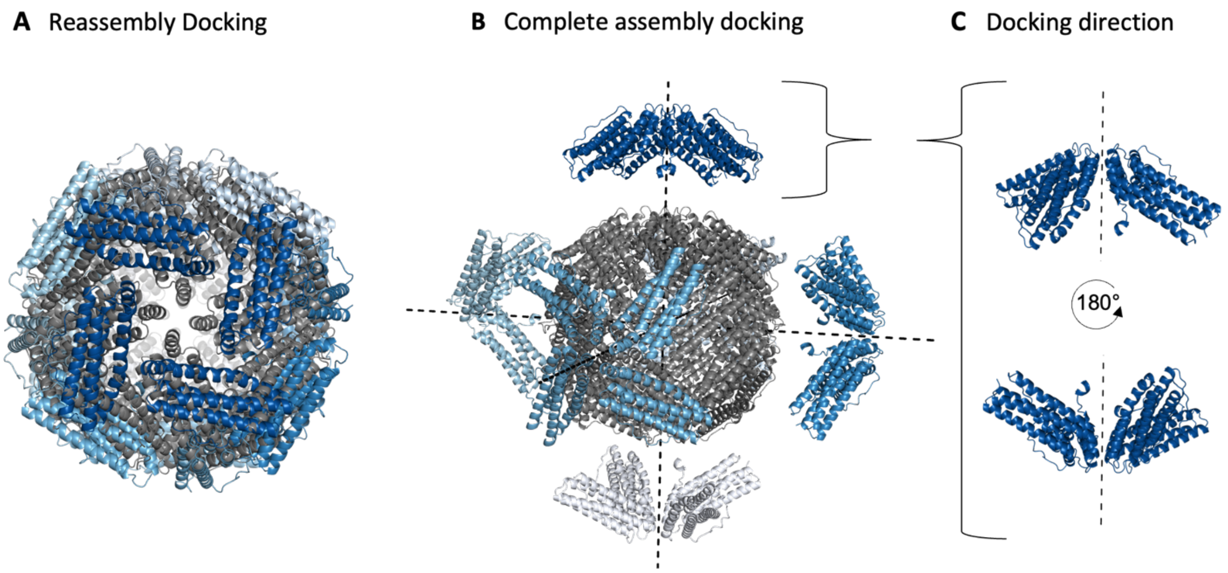


**Fig. S3: Starting conditions for local and global docking. A:** Example of the starting condition in a *Reassembly* docking scenario. The model (blue) is perturbed slightly away from the native (grey). **B:** Example of the starting condition in *Complete assembly* docking scenario. The model (blue) is perturbed far away from the native (grey)**.** Certain parameters are constrained based on the average values of $\psi, \Theta, \varphi$ and $x$ from the AFM predictions. **C:** Two docking orientations separated by a 180 degree turn perpendicular to symmetry axis are possible for Complete assembly docking.

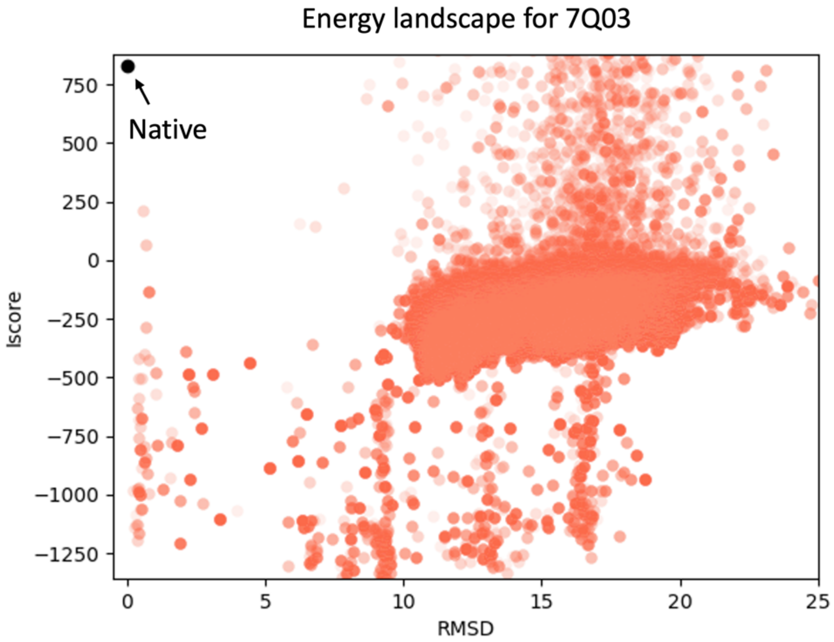


Figure 4: **Energy landscape for 7Q03 for EvoDOCK and the native structure.** EvoDOCK cannot find the native state as the Interface score (Iscore) is very high (black dot, top corner) which indicates that this is a problem with the score function (REF2015[2]) used.

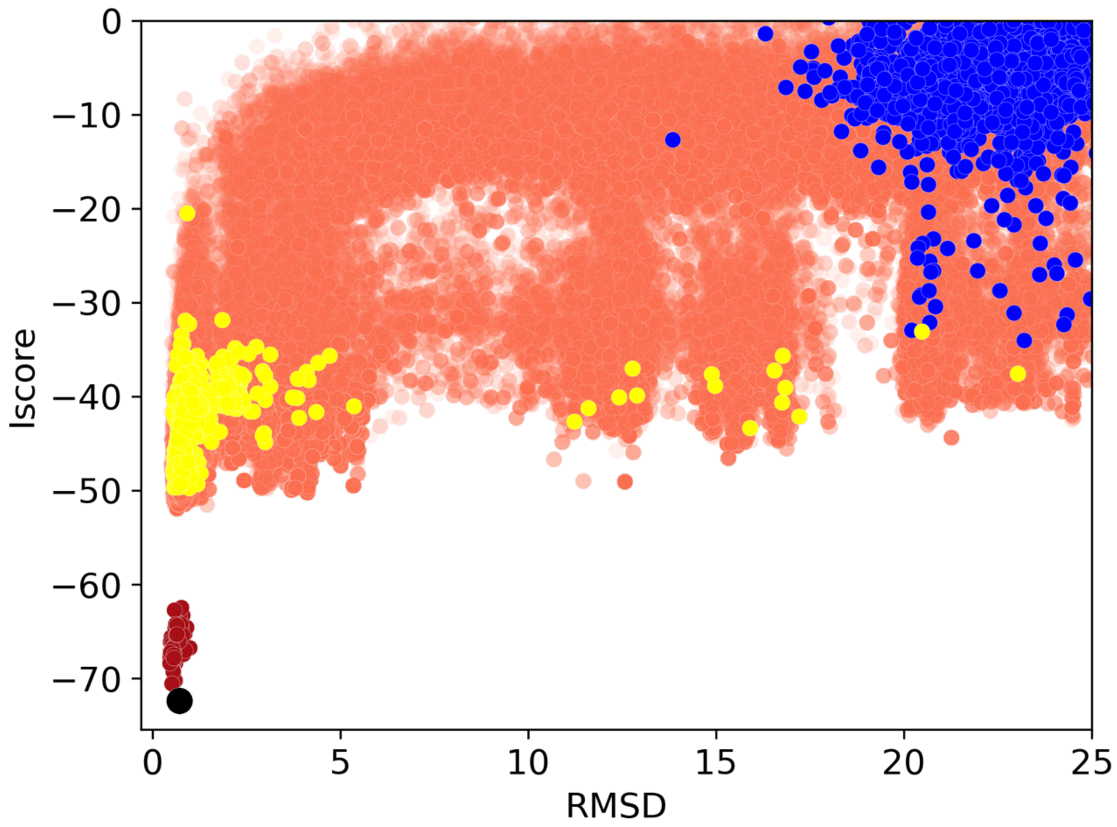


**Figure 5: Energy landscape for Complete assembly docking of 2CC9 with incorrect docking orientation.** The entire population were split up into 2 halfs were the first half was docked in the correct docking orientation and the latter not. Light orange is the full energy landscape of the entire population, dark red the relaxed models and the black dot the best energy model. The initial energy landscape of the part of the population that were docked in the wrong orientation is shown in blue. In the last generation (50) the same group is shown in yellow and it's clear that this group has learned the correct docking orientation.

**Table S1: PDBs in the benchmark and the symmetry type used for each experiment**.

| PDB | Symmetry | Reassembly symmetry type | Complete assembly symmetry type |
|---|---|---|---|
| 4DCL | T | 3 | 3 |
| 2CC9 | T | 3 | 3 |
| 3LEO | T | 3 | 3 |
| 2QQY | T | 3 | 2 |
| 7Q03 | T | 3 | 2 |
| 6M8V | T | 3 | 2 |
| 6HSB | T | 3 | 3 |
| 3WIS | O | 4 | 3 |
| 5H46 | O | 4 | 2 |
| 5EKW | O | 4 | 3 |
| 3N1I | O | 4 | 4 |
| 6H05 | O | 4 | 3 |
| 7O63 | O | 4 | 2 |
| 7OHF | O | 4 | 2 |
| 1HQK | I | 5 | 5 |
| 1T0T | I | 5 | 5 |
| 1X36 | I | 5 | 5 |
| 7B3Y | I | 5 | 3 |
| 4V4M | I | 5 | 5 |
| 1JH5 | I | 5 | 3 |
| 6ZLO | I | 5 | 3 |

**Table S2: All metric values for the Reassembly simulations**

| PDB | TM-score | | Pairwise DockQ score | | RMSD | |
|---|---|---|---|---|---|---|
| | Best Ranked | Best Cluster | Best Ranked | Best Cluster | Best Ranked | Best Cluster |
| 4DCL | 0.97 | 0.99 | 0.87 | 0.87 | 2.31 | 1.13 |
| 2CC9 | 1.00 | 1.00 | 0.86 | 0.88 | 0.61 | 0.43 |
| 3LEO | 0.93 | 0.97 | 0.57 | 0.74 | 6.00 | 2.17 |
| 2QQY | 0.96 | 0.96 | 0.86 | 0.86 | 0.83 | 0.83 |
| 7Q03 | 0.99 | 0.99 | 0.82 | 0.82 | 1.36 | 1.36 |
| 6M8V | 1.00 | 1.00 | 0.81 | 0.81 | 0.86 | 0.86 |
| 6HSB | 0.99 | 0.99 | 0.84 | 0.87 | 1.65 | 1.50 |
| 3WIS | 0.98 | 0.98 | 0.57 | 0.60 | 2.71 | 2.70 |
| 5H46 | 0.93 | 0.93 | 0.55 | 0.56 | 4.82 | 4.71 |
| 5EKW | 0.99 | 1.00 | 0.69 | 0.86 | 1.25 | 1.12 |
| 3N1I | 0.99 | 0.99 | 0.63 | 0.73 | 1.36 | 1.36 |
| 6H05 | 0.96 | 0.96 | 0.40 | 0.43 | 3.69 | 3.69 |
| 7O63 | 1.00 | 1.00 | 0.73 | 0.73 | 0.88 | 0.66 |
| 7OHF | 0.98 | 1.00 | 0.66 | 0.84 | 2.05 | 1.06 |
| 1HQK | 0.99 | 1.00 | 0.70 | 0.83 | 1.47 | 0.72 |
| 1T0T | 1.00 | 1.00 | 0.79 | 0.92 | 1.48 | 1.48 |
| 1X36 | 1.00 | 1.00 | 0.82 | 0.82 | 1.17 | 1.17 |
| 7B3Y | 1.00 | 1.00 | 0.92 | 0.93 | 0.94 | 0.62 |
| 4V4M | 0.99 | 0.99 | 0.70 | 0.70 | 1.87 | 1.87 |
| 1JH5 | 1.00 | 1.00 | 0.91 | 0.91 | 0.69 | 0.69 |
| 6ZLO | 0.99 | 0.99 | 0.76 | 0.76 | 1.97 | 1.97 |

**Table S3: All metric values for the Complete assembly simulations**

| PDB | TM-score | | Pairwise DockQ score | | RMSD | |
|---|---|---|---|---|---|---|
| | Best Ranked | Best Cluster | Best Ranked | Best Cluster | Best Ranked | Best Cluster |
| 4DCL | 0.99 | 0.99 | 0.71 | 0.81 | 1.50 | 1.50 |
| 2CC9 | 1.00 | 1.00 | 0.87 | 0.88 | 0.59 | 0.54 |
| 3LEO | 0.47 | 0.81 | 0.65 | 0.71 | 14.92 | 7.00 |
| 2QQY | 0.71 | 0.89 | 0.55 | 0.55 | 9.13 | 3.70 |
| 7Q03 | 0.99 | 1.00 | 0.81 | 0.85 | 33.64 | 1.23 |
| 6M8V | 1.00 | 1.00 | 0.62 | 0.89 | 0.83 | 0.80 |
| 6HSB | 0.99 | 0.99 | 0.85 | 0.85 | 1.71 | 1.50 |
| 3WIS | 0.49 | 0.99 | 0.39 | 0.63 | 15.88 | 2.42 |
| 5H46 | 0.91 | 0.95 | 0.53 | 0.55 | 5.36 | 3.61 |
| 5EKW | 1.00 | 1.00 | 0.73 | 0.79 | 1.26 | 0.67 |
| 3N1I | 0.99 | 0.99 | 0.69 | 0.71 | 1.84 | 1.76 |
| 6H05 | 0.94 | 0.96 | 0.39 | 0.43 | 4.66 | 3.54 |
| 7O63 | 0.99 | 1.00 | 0.72 | 0.84 | 1.32 | 0.85 |
| 7OHF | 0.99 | 0.99 | 0.79 | 0.79 | 1.46 | 1.46 |
| 1HQK | 0.99 | 1.00 | 0.64 | 0.81 | 1.90 | 0.73 |
| 1T0T | 1.00 | 1.00 | 0.80 | 0.80 | 1.35 | 1.35 |
| 1X36 | 1.00 | 1.00 | 0.83 | 0.83 | 1.11 | 1.03 |
| 7B3Y | 1.00 | 1.00 | 0.92 | 0.93 | 0.84 | 0.84 |
| 4V4M | 0.99 | 0.99 | 0.53 | 0.75 | 2.44 | 1.92 |
| 1JH5 | 1.00 | 1.00 | 0.84 | 0.85 | 0.87 | 0.84 |
| 6ZLO | 0.99 | 0.99 | 0.66 | 0.66 | 2.04 | 2.04 |

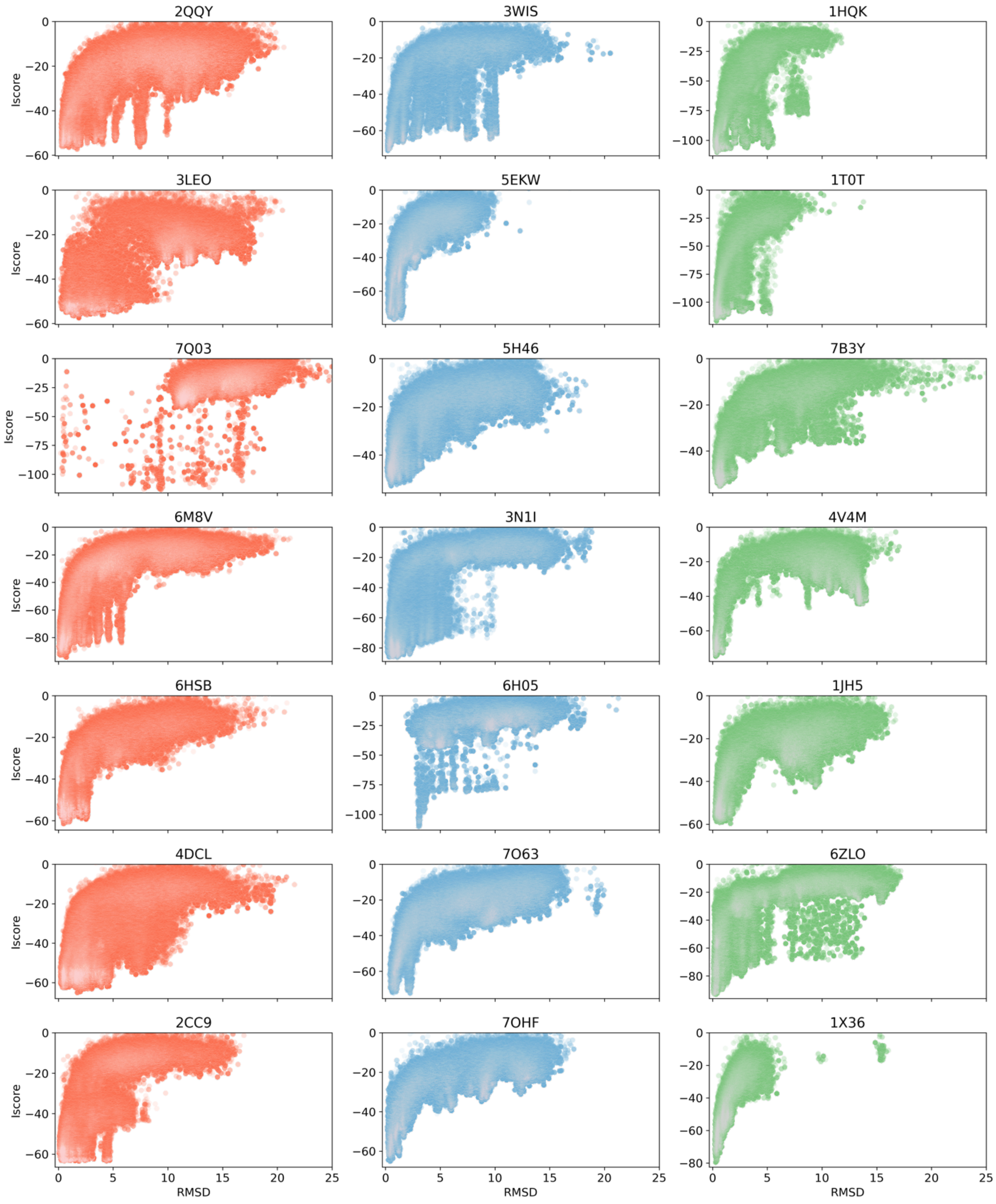


**Figure S6: Energy landscapes for Reassembly docking using native backbones.**

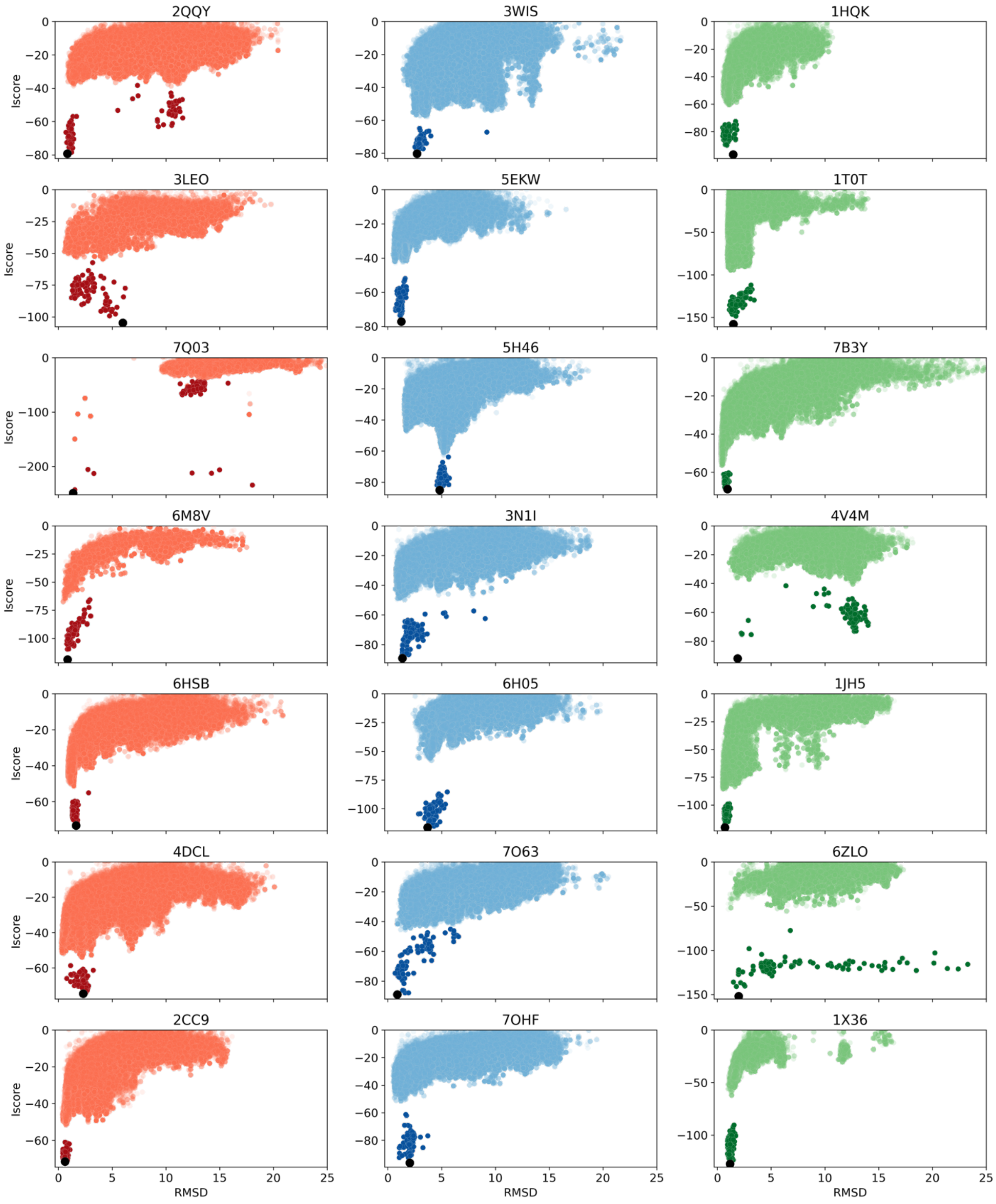


**Figure S7: Energy landscapes for Reassembly docking using AF/AFM ensemble.**

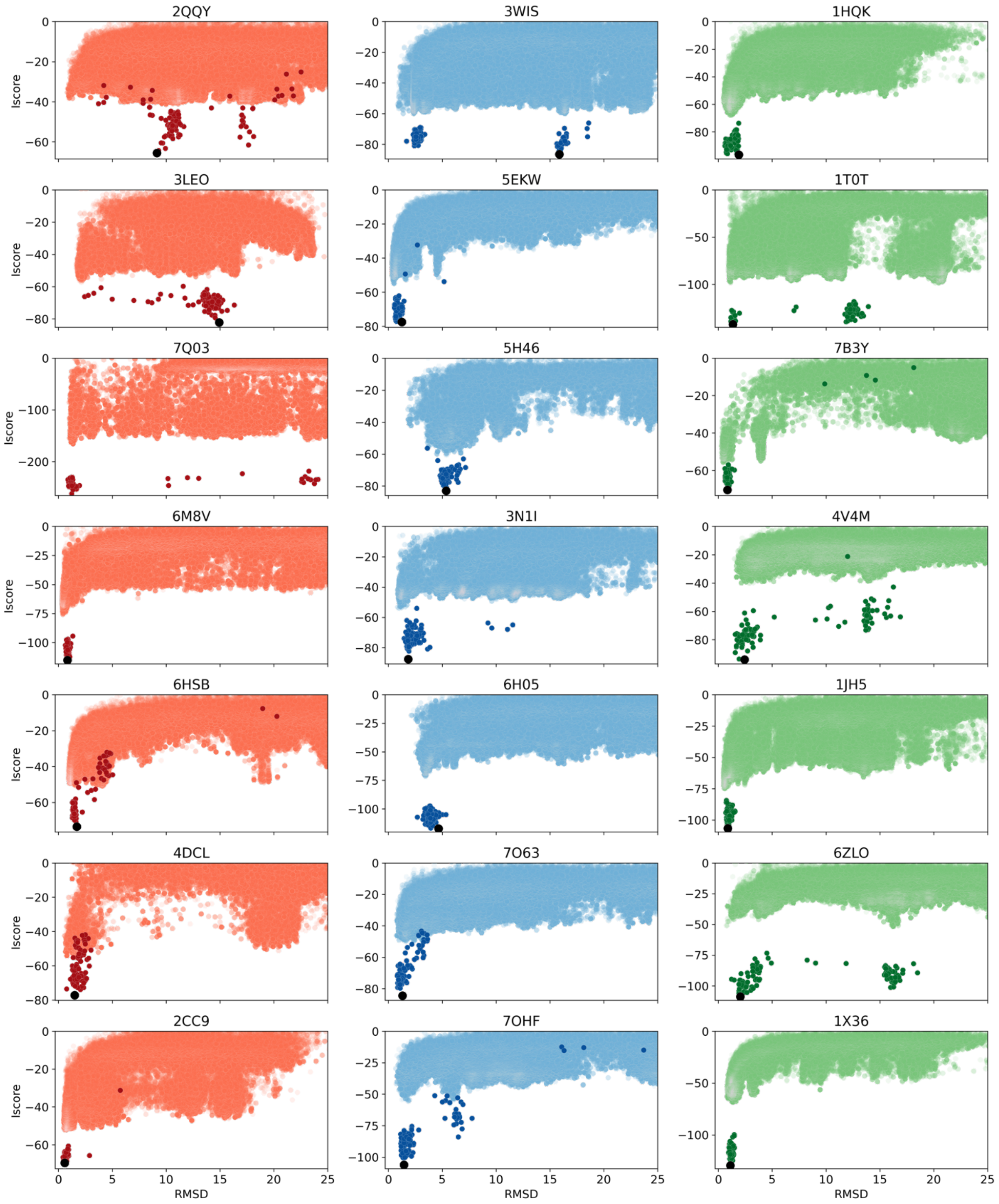


**Figure S8: Energy landscapes for Reassembly docking using AF/AFM ensemble.**

1. Berman HM*, et al.* The Protein Data Bank. *Nucleic Acids Res* **28**, 235-242 (2000).

2. Park H*, et al.* Simultaneous Optimization of Biomolecular Energy Functions on Features from Small Molecules and Macromolecules. *Journal of Chemical Theory and Computation* **12**, 6201-6212 (2016).
